# Metabolic changes of the host-pathogen environment in a *Cryptosporidium* infection

**DOI:** 10.1101/145979

**Authors:** Christopher N. Miller, Charalampos G. Panagos, Martin Kváč, Mark J. Howard, Anastasios D. Tsaousis

## Abstract

*Cryptosporidium* is an important gut microbe whose contributions towards infant and immunocompromise patient mortality rates are steadily increasing. Current techniques for diagnosing, curing or simply understanding the biology of the parasite are few and far between, relying on a combination of *in-silico* predictions modelled on a varied and unique group of organisms and medical reports. The development of an *in-vitro* culture system, using COLO-680N cells, has provided the *Cryptosporidium* community with the opportunity to expand its toolkit for investigating this disease. One area in particular that is sorely overlooked is the metabolic alterations upon infection. Existing research is extremely limited and has already shown that significant variation can be found between the metabolome of different infected host species. Using a ^1^H Nuclear Magnetic Resonance approach to metabolomics, we have explored the nature of the mouse gut metabolome as well as providing the first insight into the metabolome of an infected cell line. Through a combination of Partial Least Squares Discriminant Analysis and predictive modelling, we exhibit new and potentially game changing insights into the effects of a *Cryptosporidium parvum* infection, while verifying the presence of known metabolic changes. Of particular note is the potential contribution of host derived taurine to the diuretic aspects of the disease previously attributed to a solely parasite based alteration of the gut environment. This practical and informative approach can spearhead our understanding of the *Cryptosporidium*-host metabolic exchange and thus provide novel targets for tackling this deadly parasite.

## Introduction

Cryptosporidiosis is a disease characterised by prolonged episodes of intense diarrhoea and is the second largest cause of diarrhoeal disease and death in infants across Africa and South Asia (Checkley et al. 2014, Kotloff et al. 2013, Striepen 2013, Wanyiri et al. 2014). It is also amongst one of the highest medically important diseases of the immunocompromised, especially HIV positive patients who are at 75-100% risk of contracting the disease depending on the geographical area (Shirley et al. 2012, Wanyiri et al 2014). The pathogens responsible are parasites belonging to the apicomplexans, the *Cryptosporidium* species, of which *C. parvum* is typically the more likely (Caccio 2005, Leoni et al. 2006, Widmer and Sullivan 2012, Wielinga et al. 2008). Infection occurs when an individual ingests the oocysts of the parasite, often swallowing a contaminated water source. Water treatment options are limited to filtering, which is generally not possible at an industrial scale and UV treatment, which is both expensive and rarely available prior to the outbreak. Failing this, treatment is typically rehydration, although one drug has been shown to be effective, the broad spectrum anti-parasitic Nitazoxanide (Doumbo et al. 1997). However, the drug is far from ideal and displays a range of undesirable side effects including cytotoxicity and nausea, as well as being limited to use in cases where the patients are immunocompetent (Domjahn et al. 2014, Hussien et al. 2013, Manjunatha et al. 2016, Sparks et al. 2015).

Until recently, a significant barrier to research into cryptosporidiosis has been the absence of a combined long-term *in vivo* culturing system and comprehensive model of host parasite interactions in addition to a heavy reliance on antibody based detection both in the scientific and the medical field (Briggs et al. 2014, Checkley et al. 2014, Domjahn et al. 2014, Girouard et al. 2006, Karanis and Aldeyarbi 2011, Leitch and He 2012, Muller and Hemphill 2013, Striepen 2013). Recent papers have attempted to rectify this by proposing improved or entirely novel techniques for culturing the parasite *ex-vivo* in tissue cultures, using the cultured cancer cells as host cells (Morada et al. 2016, Muller and Hemphill 2013). A recent study identified that infections of COLO-680N cell cultures produced a longer term and higher production volume culture of the parasite compared to previously existing *in-vitro* cultures (Miller et al. 2017). These advances have allowed higher in depth microscopy-based studies and even promise to provide a solution to developing a genetic engineering platform for the parasite. However, beyond microscopy and localisation studies, the knowledgebase of the host parasite interaction remains largely undeveloped (Manjunatha, et al. 2015, Sponseller et al. 2014, Striepen 2013, Wilhelm and Yarovinsky 2014).

One area lacking study is metabolomics. Only two peer-reviewed publications have explored the concept of the infection metabolome, one on mice and the other on human faecal samples, both showing a clear relation between infection and change in metabolite levels (Ng Hublin et al. 2013, Ng Hublin et al. 2012). While working on different sample sources, each identified the hexadecanoic acid as a significant contributor to the change in the metabolome during infection. Previous studies noticed a number of metabolites, mainly amino acids, decreased in relative abundance in infected mice faeces compared to an increase seen previously in humans (Ng Hublin et al. 2012). This was explained to be most likely due to the inherent variation between the different host species metabolomes, as highlighted by Saric et al. in 2008 and highlights a pressing need for further and wider reaching studies into the metabolome of *Cryptosporidium* infections as well as the development and application of different techniques beyond the Gas Chromatography Mass Spectrometry (GC-MS) used in those papers (Ng Hublin et al. 2013, 2012, Saric et al. 2008).

^1^H Nuclear Magnetic Resonance (NMR) metabolomics is a powerful alternative to GC-MS for metabolic screening. ^1^H NMR is a simpler method that allows for a comparatively lossless analysis of metabolites, with fewer steps between sample recovery and analysis (Bezabeh et al. 2009, Hong et al. 2010, Jacobs et al. 2008, Saric, et al. 2008, Wu et al. 2010). This translates to a more reliable result in terms of quantification and reproducibility. As such, NMR has already seen use in analysing the profile of *Plasmodium falciparum*, although the metabolome of the apicomplexan parasite as a whole is almost entirely unexplored (Sengupta et al. 2016). Here, we show a novel method of analysing cryptosporidiosis-induced changes in infected mice guts metabolomes, using a ^1^H NMR approach. In addition, we have applied the same NMR based methodology to the *in vitro* infected COLO-680N cell cultures, in order to explore the similarities and differences displayed between *in-vivo* and *in-vitro* models and identify potential cross-species markers of infection.

## Materials and Methods

### Cryptosporidium

Three isolates of *C. parvum* were used in this study. The reference strain *C. parvum* Iowa II was obtained from Bunch Grass Farm in the United States, isolated from infected calves. The human isolate *C. parvum* Weru strain was supplied courtesy of Dr Martin Kváč of the Institute of Parasitology Biology Centre CAS, Czech Republic. The Weru strain was originally isolated from an infected human patient and subsequently maintained by passing through SCID mice. The final isolate used was the human isolate of *C. hominis*, supplied courtesy of Prof. Rachel Chalmers from the *Cryptosporidium* Reference Unit, Singleton Hospital of NHS Wales.

### Tissue culture

75 cm^2^ monolayers of COLO-680N were infected and maintained as per the protocols outlined in Miller *et al.* 2016, using all three isolates of *Cryptosporidium*. A control group was also established, following the same protocols as the infections, absent oocysts.

### Animals and infection

For this study, seven day old BALB/c mice were infected at the Institute of Parasitology, Biology Centre CAS using pre-established protocols detailed in Meloni and Thompson, totalling three mice per condition (Meloni and Thompson 1996). Three separate groups were used, one infected with 100,000 oocysts of *C.* parvum Iowa II, another group was infected with 100,000 oocysts of the *C. parvum* Weru isolate and the final group were given a PBS control. The groups were kept physically spate and never allowed to interact. Infection was monitored from Day-1 post-infection by aniline-carbol-methyl violet staining of faecal smears staining of faecal smears, in addition to an antigen based strip test (Milacek and Vitovec 1985), RIDA®QUICK Cryptosporidium, supplied by R-Biopharm. At ten days post-infection, the mice were euthanized by cervical dislocation and decapitation. This study was carried out in accordance with Act No 246/1992 Coll. of the Czech Republic. The protocol was approved by the Committee for Animal Welfare of Biology Centre Czech Academy of Science and the veterinary administration authorities with regards to the animal experiments.

### Sample preparation for NMR

Animal samples were retrieved from the contents of the ileum and surrounding intestinal structure by dissecting out the area of interest and washing through with three ml 100% ethanol at room temperature via syringe inserted into the opening, collecting the wash through.

Collected samples were then centrifuged for three minutes at 10,000 *g*, the supernatant discarded and the pellet weights recorded. The samples were then suspended by vortex in two ml of 75 % ethanol then transferred to a new tube and an additional five ml of 75% ethanol added.

Two ml of two mm diameter glass beads were added to the samples and agitated by vortex for 30 seconds before incubating the samples for three minutes at 80°C. The samples were vortexed for a further 30 seconds or until the sample was completely homogenised. Tissue culture samples were collected by draining the media, adding six ml of ethanol at 80°C to the culture flask and scraping the cells off the surface by cell scraper, decanting the mixture of lysed cells into 15 ml polyethylene tubes.

The samples were then decanted into two ml tubes, retaining the glass beads in the falcon tubes. The beads were washed with an additional two ml of 80°C, 75% ethanol and again the liquid was decanted into sterile two ml tubes, retaining the glass beads in the tube.

Cell debris and general detritus were removed from the samples by centrifugation at 16,000 *g* for 10 minutes and the supernatant transferred to new, sterile two ml microcentrifuge tubes. The samples were then dried via Rotorvac overnight at 40°C, suspended in 330 µl double distilled water and centrifuged at 2,500 *g* for 10 minutes. The supernatant for the samples were recombined into a single 1.5 ml microcentrifuge tube per original samples and frozen at -20 °C until the day before NMR analysis. Twenty-four hours prior to analysis, the sample tubes were placed into a freeze drier until completely desiccated. For NMR analysis, the samples were suspended in one ml of deuterated water and spiked with the sodium salt of the calibration control compound 3-(Trimethylsilyl)-1-propanesulfonic acid (DSS) to a final concentration of 20 mM and a tested pH of 7.5.

### NMR protocol and analysis

Samples were analysed using a 4-channel Bruker Avance III 14.1 T NMR spectrometer (600 MHz ^1^H) equipped with a 5 mm QCI-F cryoprobe. For controls: six separate, uninfected 25 cm^2^ COLO-680N 100% confluent monolayer cultures were analysed in addition to three uninfected BALB/c mice. Infected samples consisted of six 25 cm^2^ COLO-680N 100% confluent monolayers in addition to three Iowa infected BALB/c and three Weru infected BALB/c mice. One dimension NMR datasets were acquired with a pulse repetition rate of 5 s over 128 scans, preceded by eight equilibrating dummy scans and suppression of the residual Deuterium Oxide solvent (HDO) resonance using presaturation. Processed NMR spectrographic datasets were produced by Topspin 3.2 and analysed using Chenomx NMR Suite version 8.2. Partial Least Squares Discriminant Analysis (PLS-DA) of the Chenomx data were generated with the freely available Microsoft Excel Add-in “multi-base 2015” by Numerical Dynamics, Japan (Mutlibase for Microsoft Excel 2015). Pathway predictions were produced by the MetaboAnalyst 3.0 web tool, using a hypergeometric test and relative-betweeness centrality against *Homo sapiens* and *Mus musculus* databases for the tissue culture and mouse models respectively (Xia et al. 2015).

### Indirect Fluorescence Assays

COLO-680N cultures were seeded onto Lab-Tek, two well, Permanox chamber slides (Sigma Aldrich, Cat No. Z734640) and allowed to reach 70% confluence before infecting, following previously published protocols (Miller et al. 2017). At seven days post infection the media was aspirated from the cultures and washed twice with 1 x PBS. Fresh, pre-warmed RPMI-1640 (Sigma Aldrich, Cat. No R8758) (1% Antibiotic/Antimycotic) containing 200 nM Thermofisher Mitotracker Red CMXRos (Molecular probes; Cat. No M7512), was added to the wells and incubated in the dark at 37°C for 45 minutes. The media was removed and replaced with further pre-warmed RPMI-1640 (1% Antibiotic/Antimycotic), containing 3.5% formaldehyde, for 15 minutes at 37°C as per the manufacturer’s protocol. The cells were then briefly permeabilised with 0.2% Triton-x100 in 1x PBS for 10 minutes, washed twice with 1x PBS and four drops of SporoGlo™ or Crypt-a-glo^TM^ (WATERBORNE, INC) added, with incubation at 37°C for a further 45 minutes. The final sample was then washed three times with PBS, dried and Fluoroshield™ with DAPI (Sigma Aldrich, Cat. No F6057) was added before applying a glass coverslip and sealing. Slides were visualised by fluorescence microscopy using an Olympus IX82 or Zeiss Elyra P1 confocal microscope.

### Electron microscopy images

Aclar disks of tissue culture were infected and prepared for EM according to the protocols detailed in Miller et al. 2017.

### Ethics

All animals involved in the experiments were treated and cared for by trained members of staff according to the standards set out by Directive 2010/63/EU regarding Legislation for the protection of animals used for scientific purposes.

## Results

### Mice faecal sample extractions

Faecal samples from infected and uninfected mice were monitored using aniline-carbol-methyl violet staining (**Figure 1**) to determine validity of the control and progress of infection. Samples from both control and infected mice were taken at ten days post infection.

**Figure 1:**
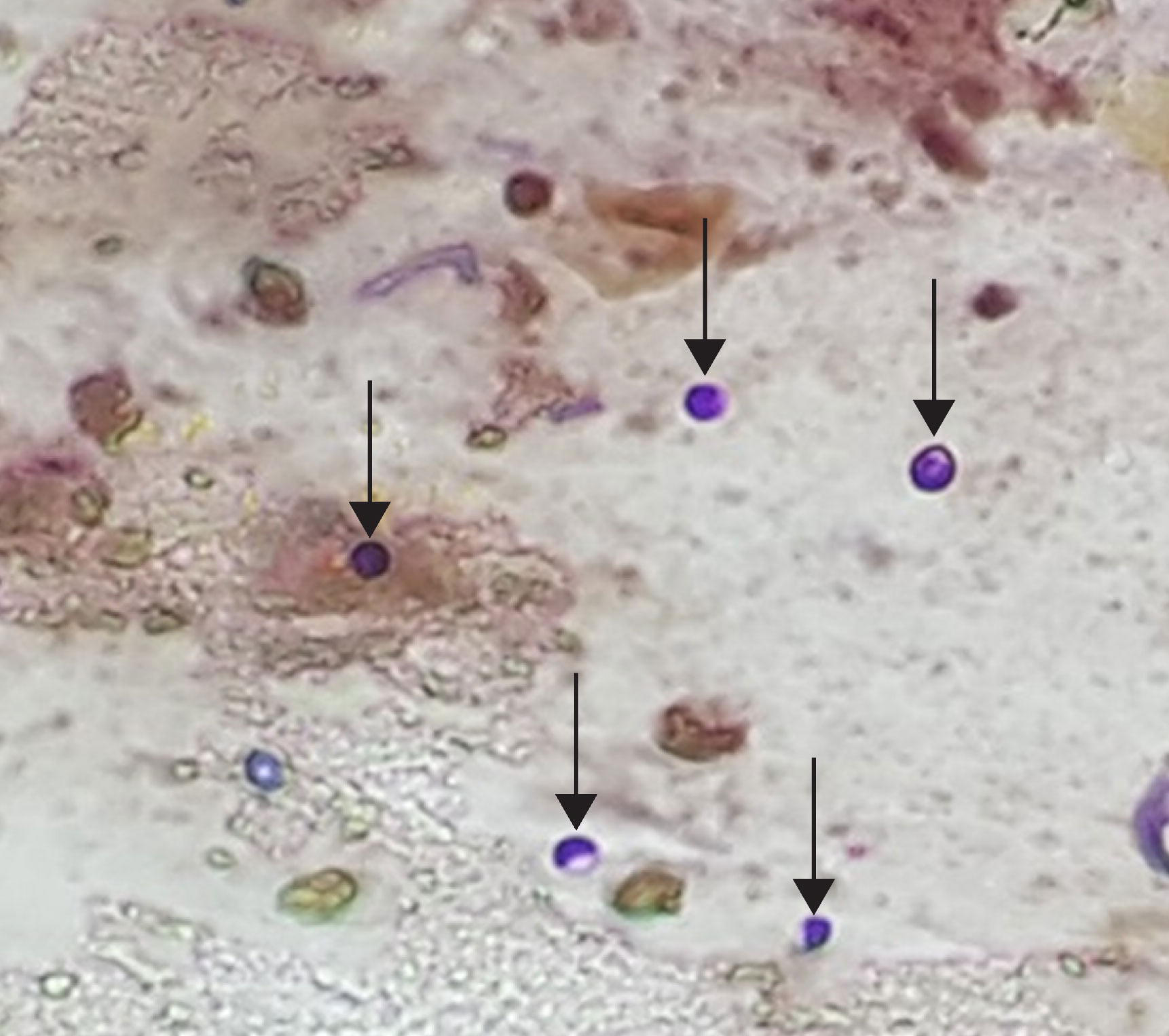
Staining of *Cryptosporidium* in faecal samples. Aniline-carbol-methyl violet stain of a faecal smear taken from a mouse in the infection group. The abundant presence of *Cryptosporidium* (arrows) indicates that the infection has been successful; and that the animal is producing oocysts.

The spectra produced by the NMR showed clear distinctions between the infected and uninfected mice, as well as distinctions between the different strains of infections (**Figure 2a**). Several metabolites were readily distinguishable prior to the metabolomics analyses, including indicators of phosphorylation; creatine and creatine phosphate (**Figure 2b**), taurine (**Figure 2c**) and lactate (**Figure 2d**). Processing the data from the mice guts (n=9) via the Chenomx Nmr Suite version 8.2 platform produced a list of 151 compounds that were extrapolated from the spectra (**Figure 3**). Statistical analysis of the data, with freely available Microsoft Excel Add-in “multi-base 2015”, by Partial Least Squares Discriminant Analysis (PLS-DA) determined significant separation of the three conditions, (uninfected control, *C. parvum* Iowa II and *C. parvum* Weru infections), whilst maintaining group cohesion (**Figure 4a**). The loading values of the variable compound contributions (**Figure 4b**), suggest certain metabolites were more significant to the separation of the groups than others. The presence of L-alanine and valine, two common amino acids, agrees with the previous literature and 2-oxoisocaproate is a component of the valine/leucine/isoleucine biosynthetic pathways reports (Ng Hublin et al. 2013, Ng Hublin et al. 2012).

**Figure 2:**
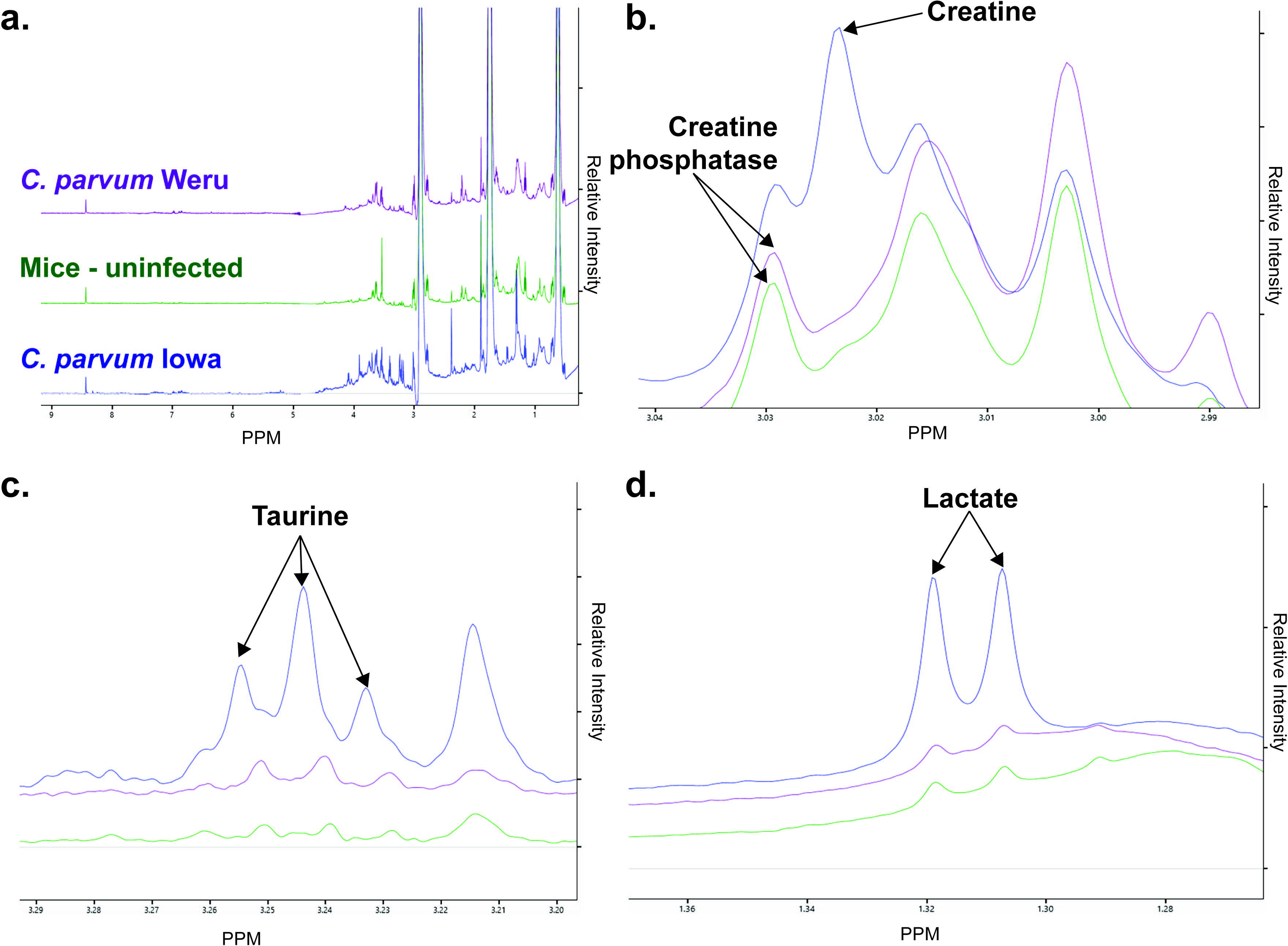
NMR Spectra of mice models of infection. **a.** Stacked NMR Spectra produced from faecal samples of the control mice (green), or either the Iowa II (blue) or Weru (purple) groups. **b.** Direct comparisons of the spectra revealed several clearly identifiably differences, including differences in creatine and creatine phosphate levels. **c.** Levels of taurine were substantially lower in the control or *C. parvum* Weru samples compared to *C. parvum* Iowa II. **d.** Lactate levels were also much higher in *C. parvum* Iowa II infected mice compared to the barely detectable levels in the control mice or *C. parvum* Weru infected groups.

**Figure 3:**
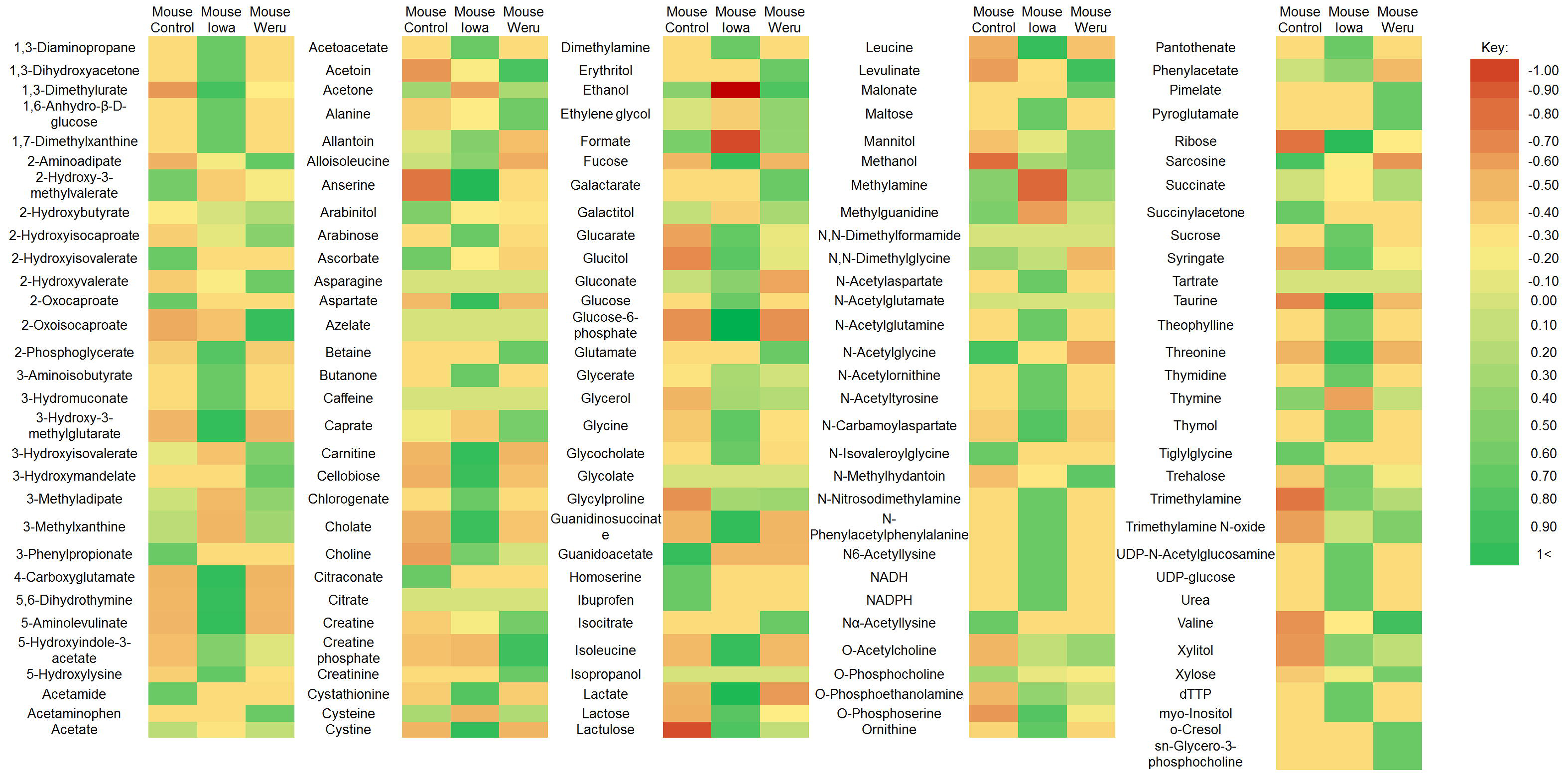
Mice Experiment Metabolites. All the metabolites identified by ^1^H NMR analysis in infected and uninfected mice were explored via PLS-DA statistical analysis. The colour coded heat map represents the significance to which each individual metabolite contributed to the identity of the sample groups, with more significant contributors tending towards green and less reliable contributors tending towards red.

**Figure 4:**
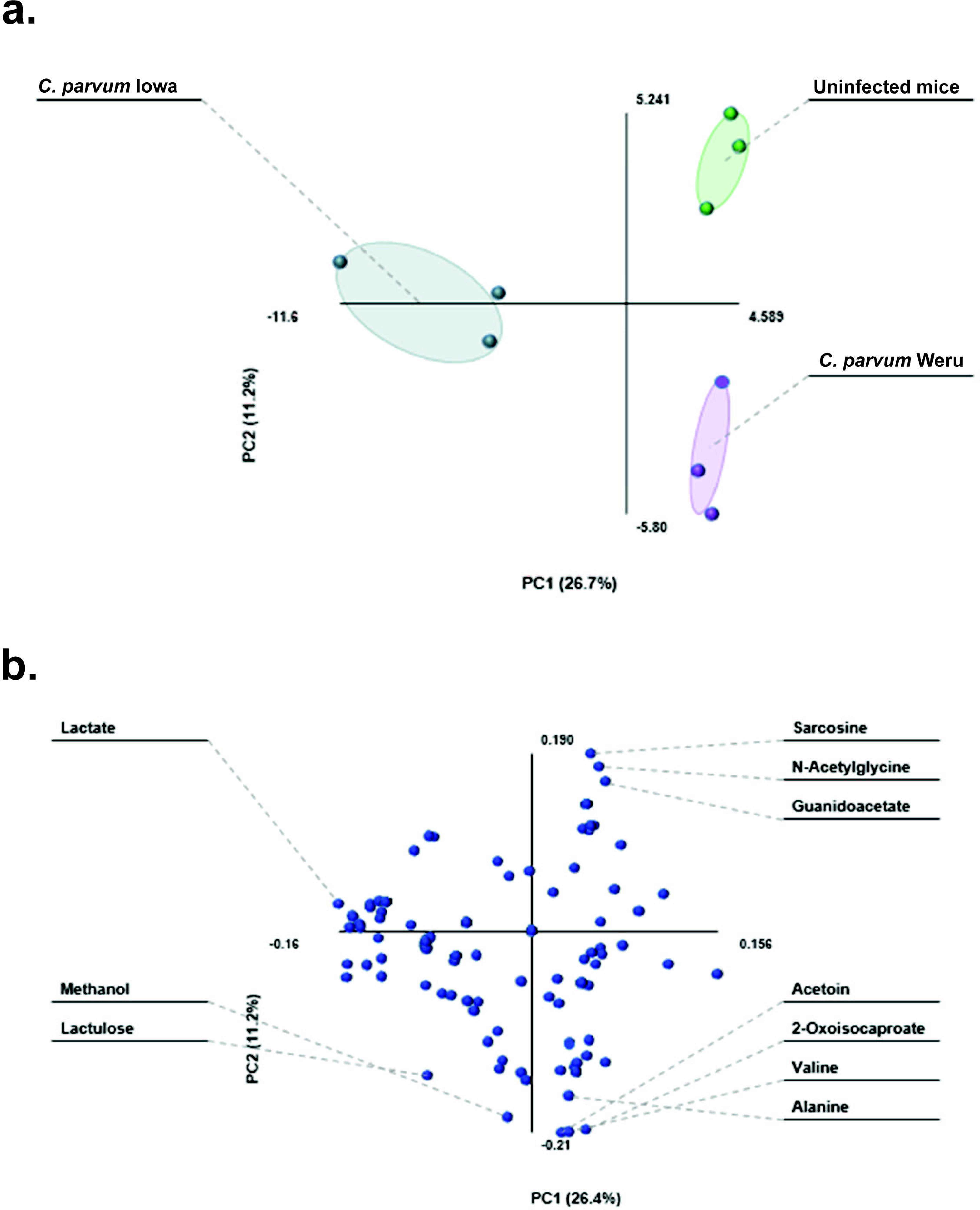
PLS-DA and loading plot of mice model NMR results. **a.** PLS-DA statistical analysis of the information provided by the Chenomx screening produced clear groupings, separating the controls (green), *C. parvum* Iowa II infections (blue) and *C. parvum* Weru infections (purple). As the grouping areas, indicated by the areas highlighted, do no overlap, it can be said that the separation between the infection conditions represent clear differences in the metabolome, which correspond to the *C. parvum* strain. **b.** The loading biplot of the PLS-DA analysis shows many of the compounds identified by Chenomx contributed towards the separation and groupings. Those on the outer most edges, for example alanine, sarcosine, lactate and lactulose, had some of the greatest influence on the amount of separation as determined by the PLS-DA.

MetaboAnalyst 3.0 based analysis of the metabolites proposed that a number of amino acid biosynthesis pathways could be altered during the course of an infection, such as the glycine, valine and taurine pathways. In addition, the mice infections displayed possible changes to other metabolic pathways (**Figure 5a**) as those pathways furthest from the x, y axis intercept, representing both the overall completeness of the pathways and number of contributing detected metabolites respectively. The pathways identified in the manner, and the compounds discovered by the NMR demonstrated that infections caused changes in at least the valine (**Figure 5c**), glycine (**Figure 5d**) and taurine amino (**Figure 5e**) acid biosynthetic pathways, in addition to several sugar pathways (**Figure 5b, f, g**).

**Figure 5:**
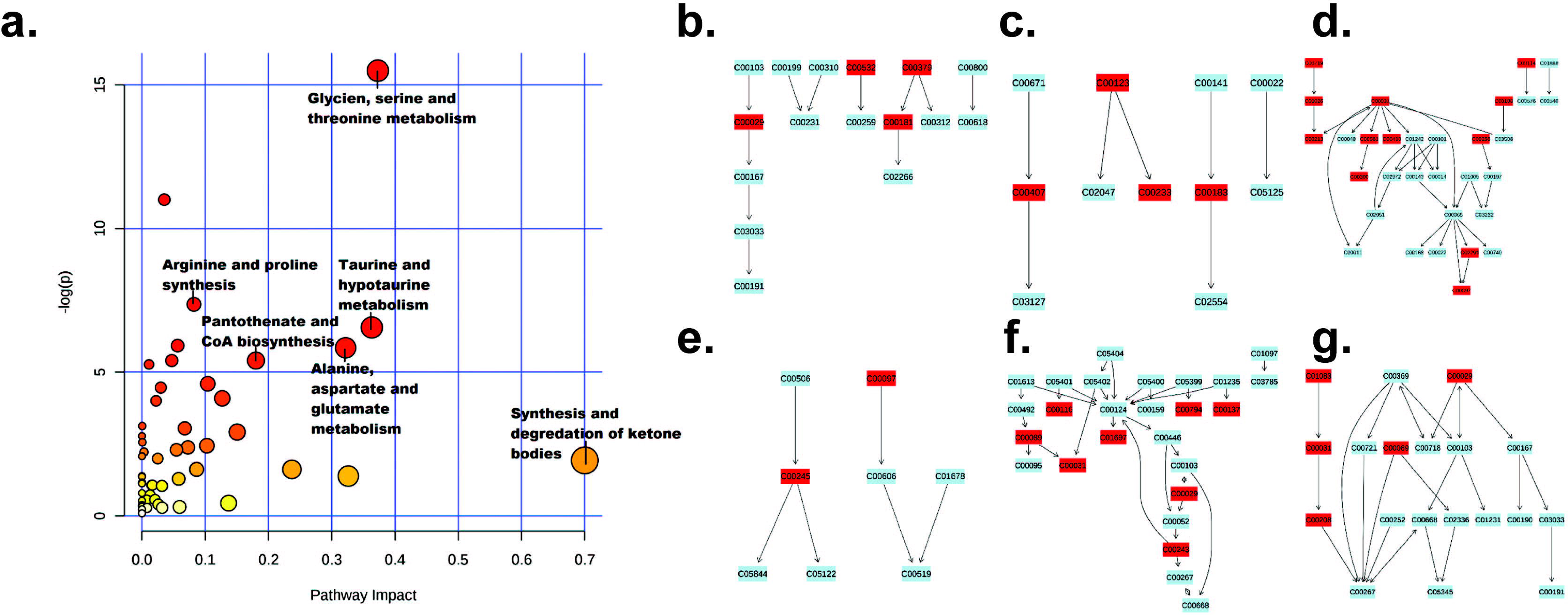
Metabolic pathways detected in mouse model NMR samples. **a.** Data analysed by MetaboAnalyst 3.0, utilising all compounds which displayed some degree of change as a result of infection, produced a graph of pathways most heavily impacted (x axis) and pathways containing the most amount of the given compounds (pathway impact: y-axis), with statistical significance of the predicted pathways increasing as the colour ranges from yellow (low) to red (high). Six pathways were chosen to be of particular interest by their position on the graph, with metabolites present in the experimental samples highlighted in red, including: **b.** pentose and glucoronate interconversions, valine, **c.** leucine and isoleucine biosynthesis, **d.** glycine serine and threonine metabolism, **e.** taurine and hypotaurine metabolism, **f.** galactose metabolism and **g.** starch and sucrose metabolism.

### Cell culture sample extractions

Extrapolated NMR data from COLO-680N (n=18) metabolite extractions, demonstrated clear differences between the each strain and species of *Cryptosporidium* used (**Figure 6**). As with the mice samples, differences between creatine, creatine phosphate, taurine and lactate (**Figures 6b**–**d**) were readily visible in the raw spectra. Chenomx analysis produced a list of 161 total compounds of varying concentrations across samples (**Figure 7**). The PLS-DA generated by the same statistical analysis as before, produced ample separation of the *Cryptosporidium-*infected and uninfected cultures, (**Figure 8a**). Furthermore, the separation of the individual infection groups suggests that differences between both *Cryptosporidium* species and within individual strains of *C. parvum*, may illicit different metabolic responses in cell cultures. The loading scores plot of the PLS-DA showed a number of amino acids contributed heavily to the separations, as well lactate, several fatty acid derivatives and taurine (**Figure 8b**).

**Figure 6:**
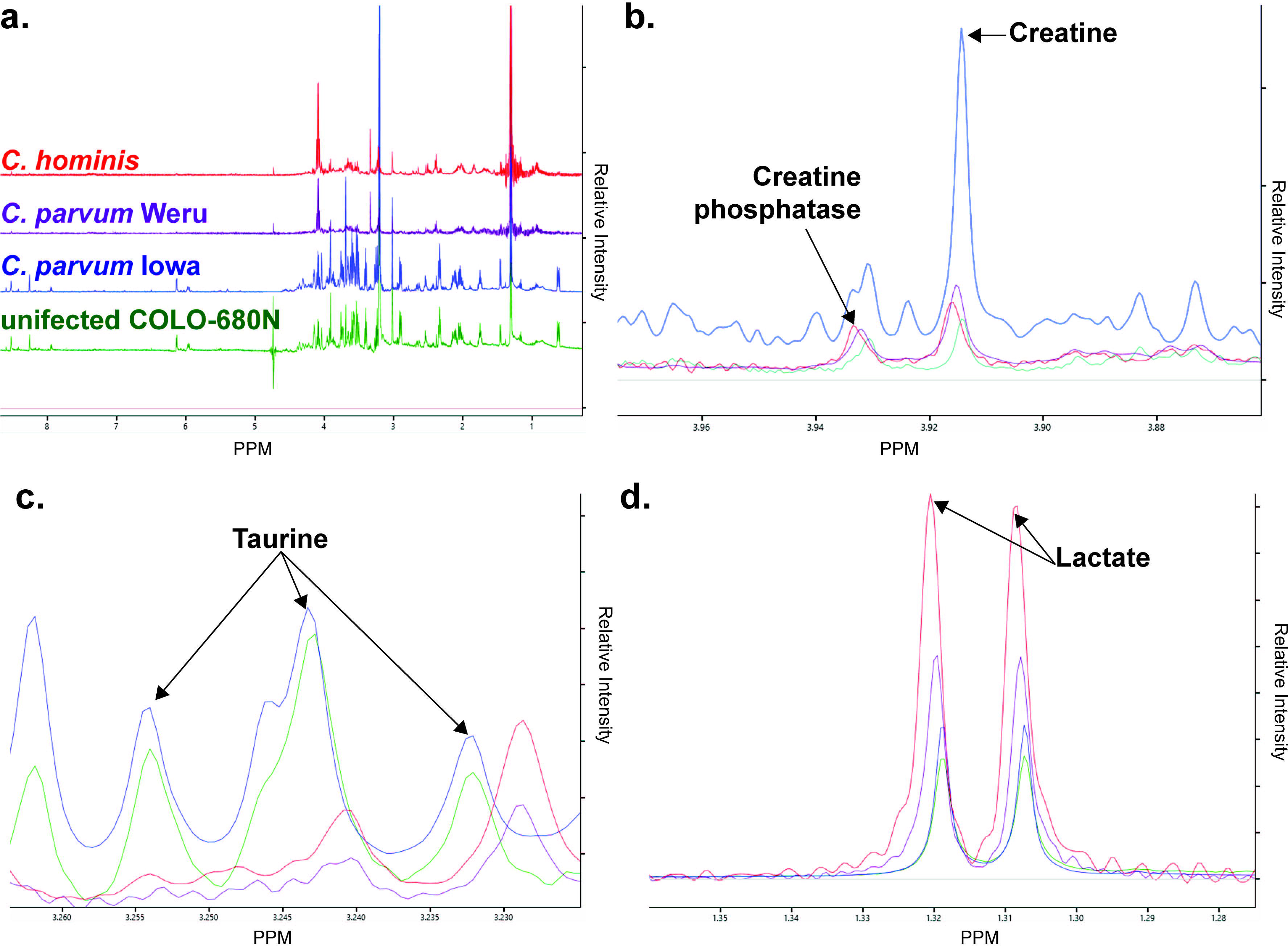
Cell Culture infection NMR spectra. **a.** Stacked NMR Spectra produced from the COLO-680N control cultures (green), or either the *C. parvum* Iowa II (blue), *C. parvum* Weru (purple), or *C. hominis* groups. Direct comparisons of the spectra revealed several clearly identifiably differences, including, again, differences in creatine and creatine phosphate (**b.**), taurine (**c.**) and lactate (**d.**) levels. Noticeably, taurine levels were almost undetectable in *C. hominis* or *C. parvum* Weru infections.

**Figure 7:**
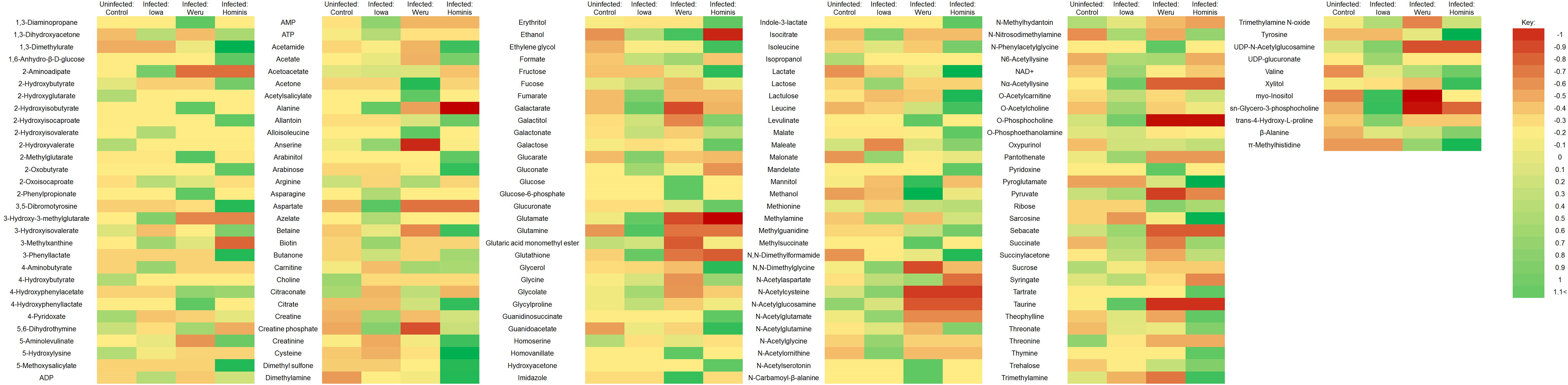
COLO-680N Experiment Metabolites. All the metabolites identified by 1H NMR analysis in infected and uninfected cells were explored via PLS-DA statistical analysis. The colour coded heat map represents the significance to which each individual metabolite contributed to the identity of the sample groups, with more significant contributors tending towards green and less reliable contributors tending towards red.

**Figure 8:**
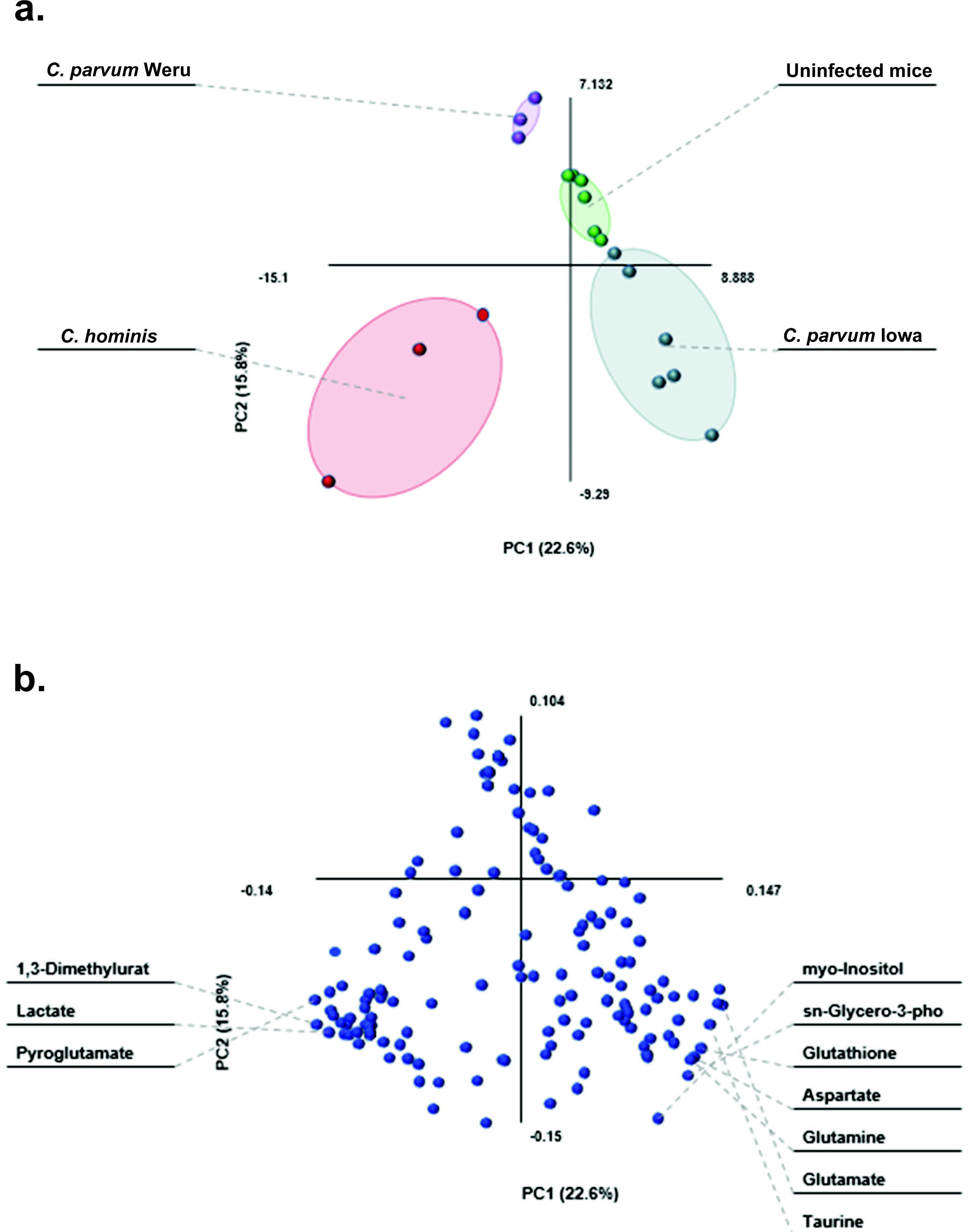
PLS-DA and loading plot of COLO-680N – infected cells NMR results. **a.** PLS-DA statistical analysis of the information provided by the Chenomx screening produced clear groupings, separating the controls (green), *C. parvum* Iowa II infections (blue), *C. parvum* Weru infections (purple) and *C. hominis* infections (red). As the grouping areas do no overlap the separation between the infection conditions again indicates that metabolome differences can be at least in part explained by different *Cryptosporidium* strains/species. **b.** The loading biplot of the PLS-DA analysis shows lactate as a significant contributor to variation, as seen before in **Figure 2b**, in addition to taurine and myo-inositol among others.

Metabolic pathway fitting via MetaboAnalyst 3.0 revealed that amino-acid biosynthesis pathways for glycine, alanine and arginine were influenced by infection. These were in addition to taurine, pantothenate and CoA biosynthetic pathways as shown in **Figure 9a**. As with **Figures 5a**–**g**, the graph shows a combination of how much of a pathway is completed by data from the NMR, as well as simply how many metabolites were detected. Among other pathways, perhaps the most significant detections were glycine (**Figure 9b**), taurine (**Figure 9c**) alanine (**Figure 9d**) and arginine (**Figure 9g**) amino acid pathways as well as, potentially the synthesis and degradation of ketones (**Figure 9e**) and pantothenate and CoA biosynthesis (**Figure 9f**).

**Figure 9:**
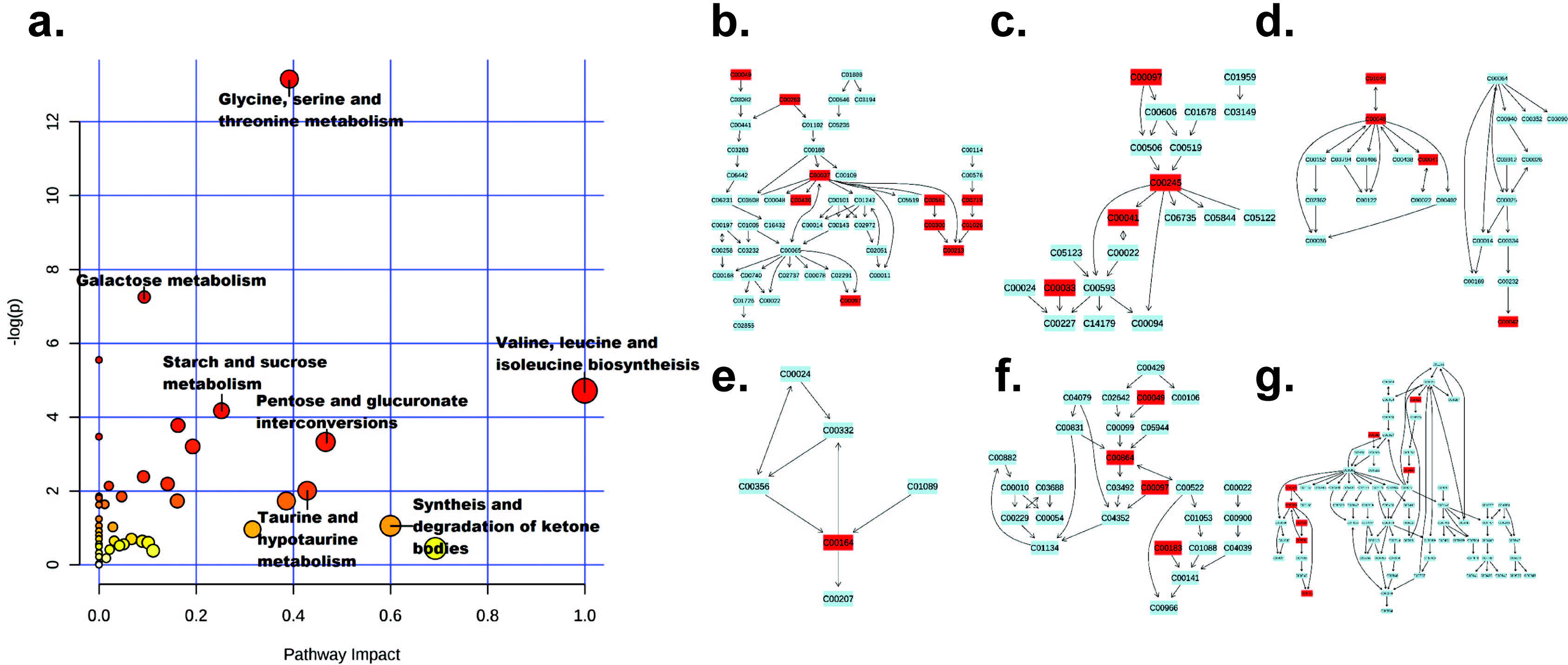
Metabolic pathways detected in cell cultures’ NMR samples. **a.** Data analysed by MetaboAnalyst 3.0, utilising all compounds which displayed some degree of change as a result of infection, produced a graph of pathways most heavily impacted (x axis) and pathways containing the most amount of the given compounds (pathway impact: y-axis), with statistical significance of the predicted pathways increasing as the colour ranges from yellow (low) to red (high). Six pathways were chosen to be of particular interest by their position on the graph, with metabolites present in the experimental samples highlighted in red, including: glycine, serine and threonine metabolism (**b.**), taurine and hypotaurine metabolism (**c.**), Alanine, aspartate and glutamate metabolism (**d.**), synthesis and degradation of ketones (**e.**), pantothenate and CoA biosynthesis (**f.**) and arginine and proline metabolism (**g.**).

### Comparison of mice faecal and COLO-680N metabolome changes

MetaboAnalyst data from **Figure 5** and **Figure 9**, demonstrate that a number of altered pathways are shared between the mice and tissue culture metabolites, particularly taurine and amino acid metabolic pathways. Taurine is involved in a number of roles, including bile acid conjugation, osmoregulation, membrane integrity and protection against oxidative free radicals. Glycine synthesis was also shown to be affected to a large degree and is involved with numerous and diverse cellular functions including purine synthesis, basic protein construction and provides the building blocks for porphyrins (Denis and Daignan-Fornier 1998, Marver et al. 1966). All of these pathways have a direct or indirect impact on the host’s mitochondrial energetic activity.

To investigate the cellular role of host mitochondria during infection, we employed an Indirect Fluorescence Assay (IFA) approach to determine if the mitochondria of the host cells were responding to *Cryptosporidium* infection (**Figure 10**). Our results demonstrate that on multiple occasions, the host mitochondria were shown to congregate in larger densities near the *Cryptosporidium* infection (**Figures 10**). Transmission Electron Microscopy images of infected cells also show abnormal host’s mitochondrial congregation around the parasitophorous vacuole (**Figure 11**).

**Figure 10:**
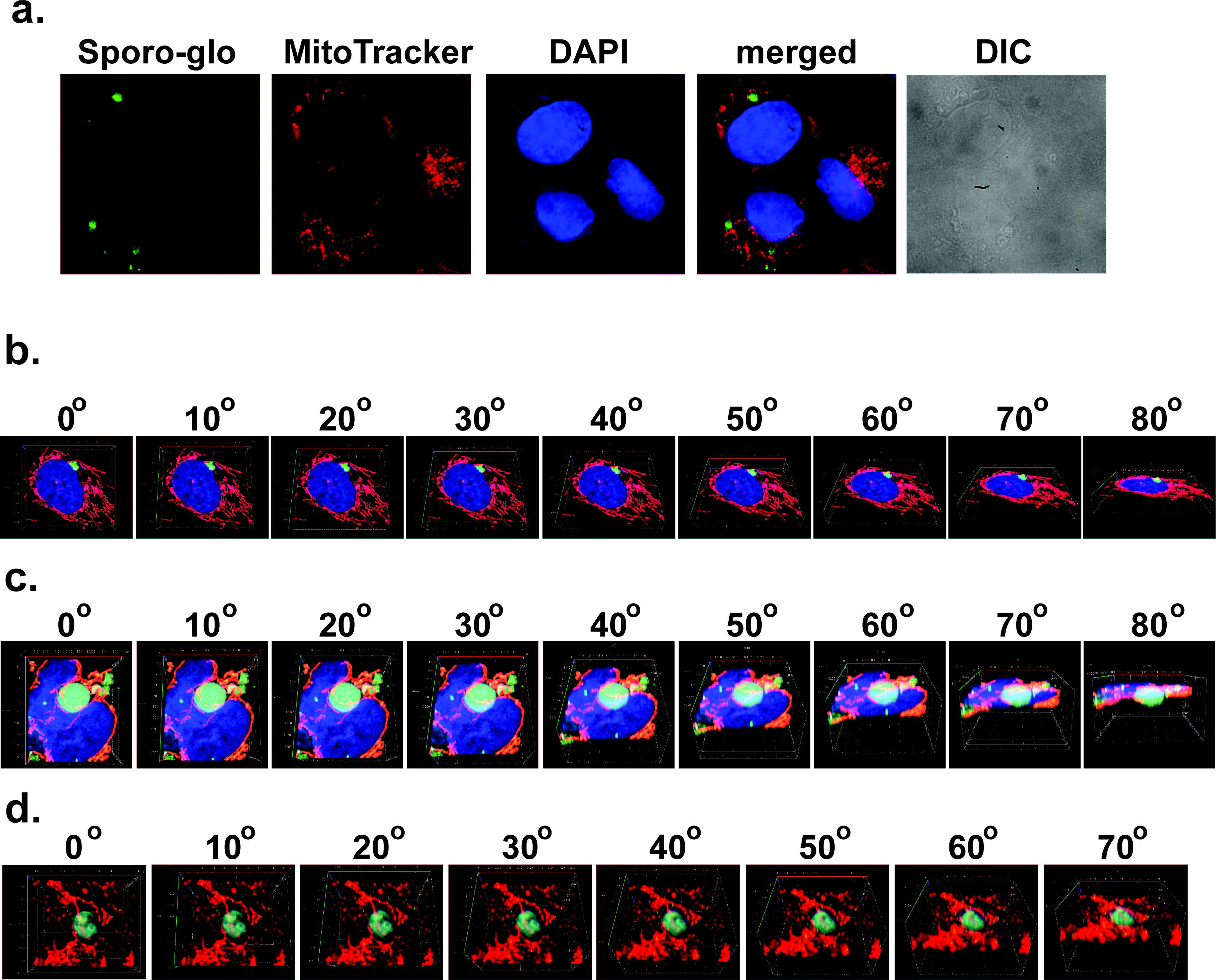
Indirect Fluorescence Assay of infected cell cultures. **a.** Fluorescence microscopy showing the staining of infected COLO-680N culture with Sporo-glo (green), MitoTracker CMXRos (red) and DAPI nuclear stain (blue). From the figure we could observe an obvious mitochondrial “clumping” and polarisation towards areas of infection, suggesting that the presence of the parasite within a host cell affects the positioning of host mitochondria. **b**. Confocal microscopy showing the localisation of Crypt-a-glo (green), MitoTracker (red) and DAPI (blue) in a 3D rendering of 31 individual, 0.16 µm thick sections, overlapping with a final representative thickness of 4.8 µm. The images are rotated around the x-axis, from 0° to 80°, showing a COLO-680N cell infected with *C. parvum* (green). Individual images of the stainings were captured in different angles, to show the infection on a three-dimensional level. A whole video showing a 360° rotation of the three-dimensional z-stack of the image is found as an animation in **Video 1**. **c**. Confocal microscopy showing the localisation of Crypt-a-glo (green), MitoTracker (red) and DAPI (blue) in a 3D rendering of 55 individual, 0.16 µm thick sections, overlapping with a final representative thickness of 8.6 µm. The images are rotated around the x-axis, from 0° to 80°, showing a COLO-680N cell infected with *C. parvum* (green). Individual images of the stainings were captured in different angles, to show the infection on a three-dimensional level. A whole video showing a 360° rotation of the three-dimensional z-stack of the image is found as an animation in **Video 2**. **d**. Confocal microscopy showing the localisation of Crypt-a-glo (green) and MitoTracker (red) in a 3D rendering of 51 individual, 0.16 µm thick sections, overlapping with a final representative thickness of 8.0 µm. The images are rotated around the x-axis, from 0° to 70°, showing mitochondria surrounding an intracellular stage of with *C. parvum* (green). Individual images of the stainings were captured in different angles, to show the infection on a three-dimensional level. A whole video showing a 360° rotation of the three-dimensional z-stack of the image is found as an animation in **Video 3**.

**Figure 11:**
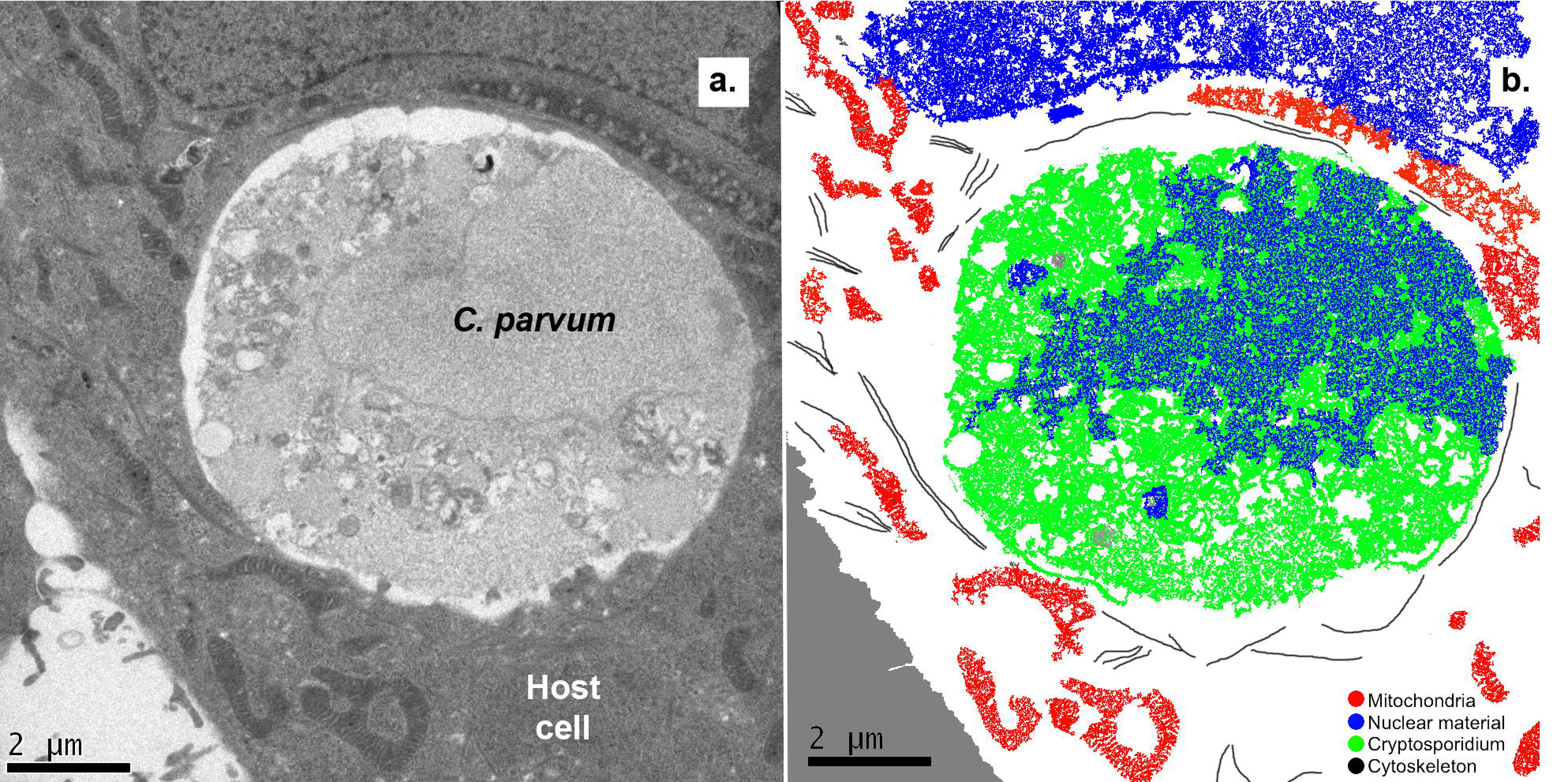
Electron microscopy of *Cryptosporidum* infected host cells. **a.** Infection of a host cell by *C. parvum*. Mitochondria of the host cell appear to closely associate with the parasitophorous vacuole surrounding the parasite, while cytoskeletal structures appear to be associated with the organelles. **b.** Cartoon of image a. demonstrating the presence of mitochondria, cytoskeleton, nuclear material and *Cryptosporidium.*

## Discussion

Solution-state ^1^H NMR offers a novel approach to metabolomics that is especially useful where sample volume sizes are particularly small (Jacobs et al 2008, Novak et al. 2006, Wu, et al. 2010). Although GC-MS holds an advantage for detecting low-levels of metabolites with unique mass signatures, for the purpose of determining the change in metabolite quantities, NMR provides a viable alternative (Bezabeh et al. 2010, Jacobs et al 2008, Saric et al. 2008, Sengupta et al. 2016, Wu et al 2010). Initial analysis of our data showed a clear distinction between the metabolic fingerprints of infected and uninfected samples, even between infections of different strains of the parasite (**Figure 4**).

Of particular importance is the degree to which these results, both from the *in-vitro* and *in-vivo*, agree with the previous literature. Our study also demonstrates that metabolic compounds L-alanine, isoleucine and succinic acid (succinate) were detected as contributors to the variance between the sample conditions that indicated infection. Moreover, even though valine was not detected in the uninfected controls, it was visible in the infected samples and in agreement with previous reports (Ng Hublin et al. 2013, Ng Hublin et al. 2012).

The predictive fits of the metabolic pathways highlighted a remarkable selection of pathways including several involved in amino acid biosynthesis, sugar metabolism, CoA biosynthesis and taurine biosynthesis. Of the predicted metabolic pathways, which have been previously shown to be influenced by infection, there are several whose presence should have been projected, such as the amino acid biosynthesis pathways for alanine and glycine as the previous reports had already highlighted their potential involvement reports (Ng Hublin et al. 2013, Ng Hublin et al. 2012).

As a parasite, *Cryptosporidium* is dependent on host derived biosynthetic pathways for survival. For example, *C. parvum* is incapable of producing the majority of amino acids *de-novo*, instead relying heavily on the import of host metabolites via active channelling (Abrahamsen et al. 2004). The biosynthetic pathway for glycine, threonine and serine was upregulated, in both cell culture and animal experimentations, with particularly high levels of glycine detected. Both *C. parvum* and *C. hominis* are incapable of manufacturing these amino acids *de novo*, instead relying on scavenging host serine and glycine, utilising serine and glycine hydroxymethyltransferases to convert one to the other when needed (Abrahamsen et al. 2004, Doyle et al. 1998). The reliance on host amino acids could provide a novel method for combating the infection, based upon previous studies that identified other amino acid metabolic chains as potential targets (Clark 1999, Doyle et al. 1998). For example, glycine reuptake inhibitors (GRIs) that are often used in treating schizophrenia, could be utilised to partially starve the parasite of the metabolite.

In addition to the amino acid biosynthesis pathways, it is also apparent that taurine synthesis is also implicated in the metabolic profile of the disease as shown in the presented analyses; taurine has frequently been used in the past as an agent for inducing excystation for *in-vitro* cultures as sodium taurochloate (Feng et al. 2006, Gold et al. 2001, Kar et al. 2011, King et al. 2012). In the host, taurine has a number of roles, those relevant to the cell types involved include: cell membrane integrity, osmoregulation and adipose tissue regulation. Previous metabolomic studies of faecal samples from *Cryptosporidium*-infected patients revealed increased taurine concentrations, explained by the characteristic decline in gut absorption as a result of villi malformation by the parasite (Goodgame et al. 1995, Kapembwa et al. 1990). However, an even greater increase in taurine levels was observed in the infected COLO-680N cell cultures, wherein malabsorption is not an applicable explanation. In addition to the pathways and the relevant metabolites featured in **Figures 5** and **9**, there were also a number of potentially important metabolites not represented. Similarly observed, was an increase in the abundance of adenosine derivatives (AMP, ADP and ATP); all showed increased abundance in infected cells and mice in *C. parvum* Iowa II infections, along with a similar increase in creatine levels in *C. parvum* Weru infections. This heavily implicates a role for the host mitochondria in the context of infection as each species and strain used lacks the creatine kinase needed to produce creatine phosphate, which typically operate in localisation with mitochondria. Levels of pyruvate in *C. hominis* cell and pantothenate in *C. parvum* Iowa II infections suggest a role for oxidatative phosphorylation. This is of particular interest as it has been established that the *C. parvum* genome contains a sequence for a potential pantothenate scavenging protein (Augagneur et al. 2013). Moreover, the further increase in lactate levels detected in *C. hominis* cell cultures and *C. parvum* Iowa II mouse infected samples, compared to the controls, indicate a strong contribution from anaerobic pathways most likely from the host. This suggests that more ATP is being produced than the oxidative capacity of the host mitochondria can maintain, producing a net increase in lactate as the oxygen debt increases. This holds particular interest as a theory of *Cryptosporidium’s* targets of parasitism include ATP production pathways, similarly to the intracellular rhizarian *Mikrocytos mackini* (Burki et al. 2013).

These data suggest that *C. parvum* and *C. hominis* infections may be directly or indirectly inducing an increase in host mitochondrial activity. If factual, this would result in a large number of oxygen free radicals being produced by the metabolic machinery. Consequently, cell(s) would respond with a matching increase in the synthesis of antioxidants such as taurine, which also sees increases during infection (Giris et al. 2008, Green et al. 1991, Zhang et al. 2004). Support for this hypothesis can be seen in the way host mitochondria appear to congregate around the *Cryptosporidium* infection (e.g. parasitophorous vacuole) (**Figures 10** and **11**). Nevertheless, taurine also plays another role within cells, for example as a diuretic. Taurine is involved in the maintenance of the ionised forms of magnesium and potassium within the cell, producing a diuretic effect that may contribute towards the characteristic water-loss of a patient with cryptosporidiosis (Kapembwa et al. 1990, Lin et al. 2016, Niggli et al. 1982, Yu et al. 2016). Furthermore, it has been found that taurine levels influence production of short chained fatty acid, another aspect of host biology theorised to be scavenged by *C. parvum* and *C. hominis* (Guo et al. 2016, Seeber and Soldati-Favre 2010, Yu et. 2016). The detection of a rise in taurine levels *in-vitro* further suggest that the increase in taurine typically detected in cryptosporidiosis patients’ stool, is more than simply the result of the guts decrease in absorptive qualities. It is likely that the intra-cellular role of taurine in this disease has been overlooked and that the pathophysiology of this disease is more complicated than currently understood, and extends beyond simple villi degradation.

Lastly, these results alone provide a promising method of determining infections via a possible comparative ^1^H NMR of patient and reference biopsies. This method offers an alternative approach in the medical field, where current methods of diagnosis are reliant on separate methods to achieve the same result as NMR, with infections detected by laborious and often inaccurate microscopy and strain typing dependant on successful PCR.

In conclusion, we have demonstrated for the first time that the use of ^1^H NMR in the context of both medical and scientific applications is indispensable in the fight against cryptosporidiosis. With the application of a more user-friendly and reproducible approach of metabolomics, through the ^1^H NMR methodology described in this paper, it will now be easier for the *Cryptosporidium* community to further explore the remaining aspects of the disease metabolome in patients’ samples. Future experiments would be best approached by increasing the number of strains analysed both *in-vitro* and *in-vivo* to test the relevant proposed hypotheses. Additionally, elucidating the more pathogenic influences of taurine biosynthesis in the pathobiology of cryptosporidiosis is critical. With these data, a metabolomics based method of diagnosing and treating the disease could become a reality.

## Abbreviations

NMR: Nuclear Magnetic Resonance
DSS: 3-(Trimethylsilyl)-1-propanesulfonic acid, sodium salt
PCA: Principal component analysis
PLS-DA: Partial Least Squares Discriminant Analysis
UV: Ultraviolet
HIV: Human Immunodeficiency Virus
GC-MS: Gas Chromatography-Mass Spectrometry
HDO: Deuterium Oxide
IFA: Indirect Fluorescence Assay
PCR: Polymerase Chain Reaction
DAPI: 4′,6-diamidino-2-phenylindole
PBS: Phosphate-buffered saline
EM: Electron microscopy
SCID: Severe Combined Immunodeficiency Disease
ATP: adenosine triphosphate
AMP: adenosine monophosphate
ADP: adenosine diphosphate
CoA: Coenzyme A
GRIs: glycine reuptake inhibitors

## Declarations

The authors have declared that the research was conducted in the absence of any commercial or financial relationships that could be construed as a potential conflict of interest

Funding was provided by the BBSRC, Wellcome Trust and a Microbiology Society Research Visit Grant.

## Acknowledgements

This research was supported by BBSRC research grant (BB/M009971/1) to Dr. Anastasios Tsaousis and a Wellcome Trust Equipment Grant 091163/Z/10/Z to Dr. Mark J. Howard. Christopher N. Miller is supported by a GTA studentship from the School of Biosciences, University of Kent and a Research Visit Grant award from the Microbiology Society. Martin Kvac is supported by the Czech Science Foundation (project No. 15-01090S). We thank Dr. Michelle Rowe for NMR technical support at Kent and members of the Dr. Tsaousis and Dr. Kvac laboratories for their intellectual and methodological support. We would also like to thank Matthew Lee and Matthew D. Badham from the University of Kent for the assistance in using the confocal microscope.

## References

Abrahamsen MS, Templeton TJ, Enomoto S, Abrahante JE, Zhu G, Lancto CA, Deng M, Liu C, Widmer G, Tzipori S, et al. 2004. Complete genome sequence of the apicomplexan, *Cryptosporidium parvum*. Science. Apr 16;304:441-445. Epub 2004/03/27.

Augagneur Y, Jaubert L, Schiavoni M, Pachikara N, Garg A, Usmani-Brown S, Wesolowski D, Zeller S, Ghosal A, Cornillot E, et al. 2013. Identification and functional analysis of the primary pantothenate transporter, PfPAT, of the human malaria parasite *Plasmodium falciparum*. J Biol Chem. Jul 12;288:20558-20567. Epub 2013/06/05.

Bezabeh T, Somorjai RL, Smith ICP. 2009. MR metabolomics of fecal extracts: applications in the study of bowel diseases. Magnetic Resonance in Chemistry. 47:S54-S61.

Briggs AD, Boxall NS, Van Santen D, Chalmers RM, McCarthy ND. 2014. Approaches to the detection of very small, common, and easily missed outbreaks that together contribute substantially to human *Cryptosporidium* infection. Epidemiol Infect. Sep;142:1869-1876. Epub 2014/04/03.

Burki F, Corradi N, Sierra R, Pawlowski J, Meyer Gary R, Abbott Cathryn L, Keeling Patrick J. 2013. Phylogenomics of the Intracellular Parasite *Mikrocytos mackini* Reveals Evidence for a Mitosome in Rhizaria. Current Biology. 8/19/;23:1541-1547.

Caccio SM. 2005. Molecular epidemiology of human cryptosporidiosis. Parassitologia. Jun;47:185-192. Epub 2005/10/29.

Checkley W, White AC, Jr., Jaganath D, Arrowood MJ, Chalmers RM, Chen XM, Fayer R, Griffiths JK, Guerrant RL, Hedstrom L, et al. 2014. A review of the global burden, novel diagnostics, therapeutics, and vaccine targets for *Cryptosporidium*. Lancet Infect Dis. Sep 29. Epub 2014/10/04.

Clark DP. 1999. New Insights into Human Cryptosporidiosis. Clinical Microbiology Reviews. 12:554-563.

Denis V, Daignan-Fornier B. 1998. Synthesis of glutamine, glycine and 10-formyl tetrahydrofolate is coregulated with purine biosynthesis in *Saccharomyces cerevisiae*. Molecular and General Genetics MGG. 259:246-255.

Domjahn BT, Hlavsa MC, Anderson B, Schulkin J, Leon J, Jones JL. 2014. A survey of U.S. obstetrician-gynecologists’ clinical and epidemiological knowledge of cryptosporidiosis in pregnancy. Zoonoses Public Health. Aug;61:356-363. Epub 2013/10/15.

Doumbo O, Rossignol JF, Pichard E, Traore HA, Dembele TM, Diakite M, Traore F, Diallo DA. 1997. Nitazoxanide in the treatment of cryptosporidial diarrhea and other intestinal parasitic infections associated with acquired immunodeficiency syndrome in tropical Africa. Am J Trop Med Hyg. Jun;56:637-639. Epub 1997/06/01.

Doyle PS, Kanaani J, Wang CC. 1998. Hypoxanthine, guanine, xanthine phosphoribosyltransferase activity in *Cryptosporidium parvum*. Exp Parasitol. May;89:9-15. Epub 1998/05/29.

Feng H, Nie W, Sheoran A, Zhang Q, Tzipori S. 2006. Bile acids enhance invasiveness of *Cryptosporidium spp*. into cultured cells. Infect Immun. Jun;74:3342-3346. Epub 2006/05/23.

Giris M, Depboylu B, Dogru-Abbasoglu S, Erbil Y, Olgac V, Alis H, Aykac-Toker G, Uysal M. 2008. Effect of taurine on oxidative stress and apoptosis-related protein expression in trinitrobenzene sulphonic acid-induced colitis. Clin Exp Immunol. 152:102.

Girouard D, Gallant J, Akiyoshi DE, Nunnari J, Tzipori S. 2006. Failure to propagate *Cryptosporidium spp*. in cell-free culture. J Parasitol. Apr;92:399-400. Epub 2006/05/30.

Gold D, Stein B, Tzipori S. 2001. The utilization of sodium taurocholate in excystation of *Cryptosporidium parvum* and infection of tissue culture. J Parasitol. Oct;87:997-1000. Epub 2001/11/07.

Goodgame RW, Kimball K, Ou C-N, White AC, Genta RM, Lifschitz CH, Chappell CL. 1995. Intestinal function and injury in acquired immunodeficiency syndrome—related cryptosporidiosis. Gastroenterology. 1995/04/01;108:1075-1082.

Green TR, Fellman JH, Eicher AL, Pratt KL. 1991. Antioxidant role and subcellular location of hypotaurine and taurine in human neutrophils. Biochim Biophys Acta. Jan 23;1073:91-97. Epub 1991/01/23.

Guo F, Zhang H, Payne HR, Zhu G. 2016. Differential Gene Expression and Protein Localization of *Cryptosporidium parvum* Fatty Acyl-CoA Synthetase Isoforms. J Eukaryot Microbiol. Mar-Apr;63:233-246. Epub 2015/09/29.

Hong YS, Ahn YT, Park JC, Lee JH, Lee H, Huh CS, Kim DH, Ryu DH, Hwang GS. 2010. 1H NMR-based metabonomic assessment of probiotic effects in a colitis mouse model. Arch Pharmacal Res. 33:1091.

Hussien SM, Abdella OH, Abu-Hashim AH, Aboshiesha GA, Taha MA, El-Shemy AS, El-Bader MM. 2013. Comparative study between the effect of nitazoxanide and paromomycine in treatment of cryptosporidiosis in hospitalized children. J Egypt Soc Parasitol. Aug;43:463-470. Epub 2013/11/23.

Jacobs DM, Deltimple N, van Velzen E, van Dorsten FA, Bingham M, Vaughan EE, van Duynhoven J. 2008. (1)H NMR metabolite profiling of feces as a tool to assess the impact of nutrition on the human microbiome. NMR Biomed. 21:615.

Kapembwa MS, Bridges C, Joseph AE, Fleming SC, Batman P, Griffin GE. 1990. Ileal and jejunal absorptive function in patients with AIDS and enterococcidial infection. J Infect. Jul;21:43-53. Epub 1990/07/01.

Kar S, Daugschies A, Cakmak A, Yilmazer N, Dittmar K, Bangoura B. 2011. *Cryptosporidium parvum* oocyst viability and behaviour of the residual body during the excystation process. Parasitol Res. Dec;109:1719-1723. Epub 2011/05/24.

Karanis P, Aldeyarbi HM. 2011. Evolution of *Cryptosporidium* in vitro culture. Int J Parasitol. Oct;41:1231-1242. Epub 2011/09/06.

King BJ, Keegan AR, Phillips R, Fanok S, Monis PT. 2012. Dissection of the hierarchy and synergism of the bile derived signal on *Cryptosporidium parvum* excystation and infectivity. Parasitology. Oct;139:1533-1546. Epub 2012/08/17.

Kotloff KL, Nataro JP, Blackwelder WC, Nasrin D, Farag TH, Panchalingam S, Wu Y, Sow SO, Sur D, Breiman RF, et al. 2013. Burden and aetiology of diarrhoeal disease in infants and young children in developing countries (the Global Enteric Multicenter Study, GEMS): a prospective, case-control study. Lancet. Jul 20;382:209-222. Epub 2013/05/18.

Leitch GJ, He Q. 2012. Cryptosporidiosis-an overview. J Biomed Res. Jan;25:1-16. Epub 2012/06/12.

Leoni F, Amar C, Nichols G, Pedraza-Diaz S, McLauchlin J. 2006. Genetic analysis of *Cryptosporidium* from 2414 humans with diarrhoea in England between 1985 and 2000. J Med Microbiol. Jun;55:703-707. Epub 2006/05/12.

Lin S, Sanders DS, Gleeson JT, Osborne C, Messham L, Kurien M. 2016. Long-term outcomes in patients diagnosed with bile-acid diarrhoea. Eur J Gastroenterol Hepatol. Feb;28:240-245. Epub 2015/12/05.

Manjunatha UH, Chao AT, Leong FJ, Diagana TT. 2016. Cryptosporidiosis Drug Discovery: Opportunities and Challenges. ACS Infect Dis. Aug 12;2:530-537. Epub 2016/09/15.

Marver HS, Tschudy DP, Perlroth MG, Collins A. 1966. Delta-aminolevulinic acid synthetase. I. Studies in liver homogenates. J Biol Chem. Jun 25;241:2803-2809. Epub 1966/06/25.

Meloni BP, Thompson RC. 1996. Simplified methods for obtaining purified oocysts from mice and for growing *Cryptosporidium parvum* in vitro. J Parasitol. Oct;82:757-762. Epub 1996/10/01.

Milacek P, Vitovec J. 1985. Differential staining of cryptosporidia by aniline-carbol-methyl violet and tartrazine in smears from feces and scrapings of intestinal mucosa. Folia Parasitol (Praha). 32:50. Epub 1985/01/01.

Morada M, Lee S, Gunther-Cummins L, Weiss LM, Widmer G, Tzipori S, Yarlett N. 2016. Continuous culture of *Cryptosporidium parvum* using hollow fiber technology. Int J Parasitol. Jan;46:21-29. Epub 2015/09/06.

Muller J, Hemphill A. 2013. In vitro culture systems for the study of apicomplexan parasites in farm animals. Int J Parasitol. Feb;43:115-124. Epub 2012/09/25.

Miller CN, Jossé L, Brown I, Blakeman B, Povey J, Yiangou L, Price M, Cinatl jr J, Xue W-F, Michaelis M, et al. 2017. A cell culture platform for C*ryptosporidium* that enables long-term cultivation and the systematic investigation of its biology. Under review, 3^rd^ revision, *Scientific Reports*, Submitted March 2017.

Mutlibase for Microsoft Excel [2015. NumericalDynamics.Com: Numerical Dynamics, Japan. Freeware add-on for microsoft Excel.

Ng Hublin JS, Ryan U, Trengove R, Maker G. 2013. Metabolomic profiling of faecal extracts from *Cryptosporidium parvum* infection in experimental mouse models. PLoS One. 8:e77803. Epub 2013/11/10.

Ng Hublin JSY, Ryan U, Trengove RD, Maker GL. 2012. Development of an untargeted metabolomics method for the analysis of human faecal samples using *Cryptosporidium*-infected samples. Molecular and Biochemical Parasitology. 10//;185:145-150.

Niggli V, Sigel E, Carafoli E. 1982. Inhibition of the purified and reconstituted calcium pump of erythrocytes by micro M levels of DIDS and NAP-taurine. FEBS Lett. Feb 22;138:164-166. Epub 1982/02/22.

Novak P, Tepes P, Fistric I, Bratos I, Gabelica V. 2006. The application of LC-NMR and LC-MS for the separation and rapid structure elucidation of an unknown impurity in 5-aminosalicylic acid. J Pharm Biomed Anal. 40:1268.

Saric J, Wang Y, Li J, Coen M, Utzinger J, Marchesi JR, Keiser J, Veselkov K, Lindon JC, Nicholson JK, et al. 2008. Species variation in the fecal metabolome gives insight into differential gastrointestinal function. J Proteome Res. Jan;7:352-360. Epub 2007/12/07.

Seeber F, Soldati-Favre D. 2010. Metabolic pathways in the apicoplast of apicomplexa. Int Rev Cell Mol Biol. 281:161-228. Epub 2010/05/13.

Sengupta A, Ghosh S, Das BK, Panda A, Tripathy R, Pied S, Ravindran B, Pathak S, Sharma S, Sonawat HM. 2016. Host metabolic responses to *Plasmodium falciparum* infections evaluated by 1H NMR metabolomics. Mol Biosyst. Aug 22. Epub 2016/08/23.

Shirley DA, Moonah SN, Kotloff KL. 2012. Burden of disease from cryptosporidiosis. Curr Opin Infect Dis. Oct;25:555-563. Epub 2012/08/22.

Sparks H, Nair G, Castellanos-Gonzalez A, White AC, Jr. 2015. Treatment of *Cryptosporidium*: What We Know, Gaps, and the Way Forward. Curr Trop Med Rep. Sep;2:181-187. Epub 2015/11/17.

Sponseller JK, Griffiths JK, Tzipori S. 2014. The evolution of respiratory Cryptosporidiosis: evidence for transmission by inhalation. Clin Microbiol Rev. Jul;27:575-586. Epub 2014/07/02.

Striepen B. 2013. Parasitic infections: Time to tackle cryptosporidiosis. Nature. Nov 14;503:189-191. Epub 2013/11/16.

Wanyiri JW, Kanyi H, Maina S, Wang DE, Steen A, Ngugi P, Kamau T, Waithera T, O’Connor R, Gachuhi K, et al. 2014. Cryptosporidiosis in HIV/AIDS patients in Kenya: clinical features, epidemiology, molecular characterization and antibody responses. Am J Trop Med Hyg. Aug;91:319-328. Epub 2014/05/29.

Widmer G, Sullivan S. 2012. Genomics and population biology of *Cryptosporidium* species. Parasite Immunol. Feb-Mar;34:61-71. Epub 2011/05/21.

Wielinga PR, de Vries A, van der Goot TH, Mank T, Mars MH, Kortbeek LM, van der Giessen JWB. 2008. Molecular epidemiology of *Cryptosporidium* in humans and cattle in The Netherlands. International Journal for Parasitology. 6//;38:809-817.

Wilhelm CL, Yarovinsky F. 2014. Apicomplexan infections in the gut. Parasite Immunol. Sep;36:409-420. Epub 2014/09/10.

Wu J, An Y, Yao J, Wang Y, Tang H. 2010. An optimized sample preparation method for NMR-based faecal metabonomic analysis. Analyst. 135:1023.

Xia J, Sinelnikov IV, Han B, Wishart DS. 2015. MetaboAnalyst 3.0–making metabolomics more meaningful. Nucleic Acids Res. Jul 1;43:W251-257.

Yu H, Guo Z, Shen S, Shan W. 2016. Effects of taurine on gut microbiota and metabolism in mice. Amino Acids. Jul;48:1601-1617. Epub 2016/03/31.

Zhang M, Izumi I, Kagamimori S, Sokejima S, Yamagami T, Liu Z, Qi B. 2004. Role of taurine supplementation to prevent exercise-induced oxidative stress in healthy young men. Amino Acids. Mar;26:203-207. Epub 2004/03/26.

